# Lens Placode Modulates Extracellular Matrix Formation During Early Eye Development

**DOI:** 10.1101/2023.11.30.569417

**Authors:** Cecília G. De Magalhães, Ales Cvekl, Ruy G. Jaeger, C. Y. Irene Yan

## Abstract

The role extracellular matrix (ECM) in multiple events of morphogenesis has been well described, little is known about its specific role in early eye development. One of the first morphogenic events in lens development is placodal thickening, which converts the presumptive lens ectoderm from cuboidal to pseudostratified epithelium. This process occurs in the anterior pre-placodal ectoderm when the optic vesicle approaches the cephalic ectoderm. Since cells and ECM have a dynamic relationship of interdependence and modulation, we hypothesized that the ECM evolves with cell shape changes during lens placode formation. This study investigates changes in optic ECM including both protein distribution deposition, extracellular gelatinase activity and gene expression patterns during early optic development using chicken and mouse models. In particular, the expression of *Timp2*, a metalloprotease inhibitor, corresponds with a decrease in gelatinase activity within the optic ECM. Furthermore, we demonstrate that optic ECM remodeling depends on BMP signaling in the placode. Together, our findings suggest that the lens placode plays an active role in remodeling the optic ECM during early eye development.

## INTRODUCTION

During embryonic development, epithelial morphogenesis plays an important role in tissue and organ formation; shaping tubes, cavities, and folds; ultimately resulting in the development of three-dimensional organs. It depends on complex cellular interactions and extracellular matrix remodeling, cell proliferation, motility, shape and adhesive properties. Thus, orchestration of these morphogenetic mechanisms depends on several precisely regulated cellular and molecular factors.

The extracellular matrix (ECM) is a major driver of morphogenesis. It plays an important role in epithelial shape and differentiation during embryonic development. The composition and physical characteristics of the ECM influence multiple aspects of cell behavior (Mouw et al., 2014; Rozario and DeSimone, 2010). An example of the importance of ECM during development is branching morphogenesis (Rozario and DeSimone, 2010). The equilibrium between branching and growth of buds requires fine control of the ECM composition in specific domains. A thick ECM is formed around the bud flanks, whereas a thinner ECM is formed at the end bud tips (Fata et al., 2004; Rozario and DeSimone, 2010; Simian et al., 2001).

During early vertebrate eye development the lens placode, its adjacent surface ectoderm and underlying optic vesicle undergo highly coordinated morphogenetic processes to form the future retina, lens and cornea (Bailey et al., 2006; Cvekl and Ashery-Padan, 2014; Gunhaga, 2011). Eye morphogenesis initiates with the evagination of the optic vesicles from the neural tube, which extend laterally and approach the surface ectoderm (stage HH10-11 in chick embryo and E9 in the mouse). At this stage, two factors play a crucial role in lens placode induction: BMP signaling and the transcription factor Pax6 (De Magalhães et al., 2021). BMP signaling determines and maintains lens placodal fate in the surface ectoderm (Sjodal et al., 2007). Additionally, the expression of Pax6 marks the lens’s fate. After optic vesicle evagination, Pax6 expression in the head surface ectoderm becomes restricted to the optic vesicle and lens placodal region (Bhattacharyya & Bronner-Fraser, 2008; De Magalhães et al., 2021). Pax6 expression in the pre-lens ectoderm is essential for progression into subsequent phases. After optic vesicle evagination, the surface ectoderm cells grow in the apical-basal axis, defining the lens placode (HH12-14 in the chick and E9.5-9.75 in the mouse embryos) (De Magalhães et al., 2021). Following their thickening, placodal cells undergo reduction of the apical surface through cytoskeletal-driven cell shape changes, leading to lens placode invagination (HH15 in the chick and E10 in the mouse embryo) (Borges et al., 2011; Chauhan et al., 2009; Chauhan et al., 2015; Chauhan et al., 2011; Chow and Lang, 2001; Jidigam et al., 2015; Lang et al., 2014; Plageman et al., 2010; Plageman et al., 2011). The underlying optic vesicle also invaginates and bends so that the distal region becomes the bilayered optic cup. In mice, this simultaneous invagination depends on lens placode-derived filopodia that connect with the optic vesicle, serving as anchors (Chauhan et al., 2009).

In addition to cell shape changes, the ECM between the two epithelia also evolves during lens placode thickening. The optic ECM between the two epithelia can be morphologically divided into three regions: two basal laminae that line the optic and lens epithelia and a common interstitial matrix. This interstitial ECM contains Collagen IV, Laminin and Fibronectin prior to and during lens development (Hilfer and Randolph, 1993; Hilfer et al., 1981; Parmigiani and McAvoy, 1984; Svoboda and O’Shea, 1987). During lens placode apical-basal growth there is an increase in glycoproteins deposition and Collagen IV and Fibronectin staining pattern changes (Hendrix and Zwaan, 1975; Huang et al., 2011; Svoboda and O’Shea, 1987). Prior to placode invagination, Fibronectin staining is punctate between the optic cup and placode (Hilfer and Randolph, 1993). After invagination, both Laminin and Fibronectin staining are less intense in the interstitial layer when compared to earlier stages (Hilfer and Randolph, 1993). Taken together, these data show that ECM evolution is tightly linked with morphogenesis of the optic region.

Indeed, interruption of lens placode morphogenesis changes the expression of ECM-associated genes. In the absence of functional *Pax6* gene -a transcription factor crucial for lens development-(Liu et al., 2006; Shaham et al., 2012; Smith et al., 2009), embryos do not develop the lens placode and the expression of several ECM genes is markedly reduced (Huang et al., 2011). This shows that the composition of optic ECM is linked to the early differentiation of lens placode cells. Conversely, lens placode morphogenesis depends on a specific composition of the ECM. Lack of Fibronectin in the optic ECM arrests lens placode morphology as a cuboidal epithelia (Huang et al., 2011).

Since the dynamics of lens placode morphology and the optic ECM seem to be interdependent, we hypothesized that the ECM evolves together with the cell shape changes during early morphogenesis of the lens placode and undergo rearrangements restricted to the optic region. In this scenario, optic ECM composition and characteristics would be regulated by optic tissues during early eye development. First, we identified dynamic changes in key ECM proteins-Laminin α1 and Fibronectin-during lens placode formation in the optic region. Inhibition of BMP signaling in the presumptive lens epithelia not only disrupted placode formation but also led to ECM abnormalities. This disruption is significant as BMP signaling plays a crucial role in patterning the pre-placodal ectoderm and subsequent stages of lens formation (Cvekl and Zhang, 2017; Faber et al., 2002; Furuta and Hogan, 1998; Lang, 2004; Sjödal et al., 2007), regulating ECM changes. We also identified that *Timp2*, a metalloprotease inhibitor, is expressed only by lens placode cells during its apical-basal growth. This expression pattern correlates spatio-temporally with a specific downregulation of metalloprotease activity in the optic ECM. Together, our results suggest that the lens placode plays an important role in determining the architecture and composition of the optic ECM during its development.

## RESULTS

### The ECM evolves during lens placode formation

Our initial goal was to analyze in detail the distribution pattern of Fibronectin (Fn) and Laminin α1 (Lama1) in the optic ECM before and after lens placode development. We detected both proteins via immunofluorescence and reconstructed a 3D image of the optic region and its surrounding tissues in chicken embryos. At stage HH11, before lens placode formation, both Fn and Lama1 have a fibrillar pattern in the ECM between the optic vesicle and the pre-placodal ectoderm (Figure 1A). In contrast, at stage HH14, when lens placode is formed, both proteins show a diffuse and punctate pattern between the thickened placode and the optic vesicle (Figure 1B). The Fn fibrillar pattern is restricted to non-placodal regions, corresponding to cells that do not undergo thickening (Figure 1B-C). Lama1 labelling remains intense in non-placodal ectodermal regions while its labelling is weak in the ECM underlying the optic ectoderm (Figure 1B). Next, we investigated whether these changes also occur during mouse lens placode formation. Indeed, Lama1 also adopted a more diffuse staining pattern under the lens placode after its appearance (Sup. Figure 1 and 2). Together, these results show, firstly, that Fn and Lama1 reorganization is specific to the lens placodal region. Further, these changes are conserved in other amniotes.

**Figure 1.**
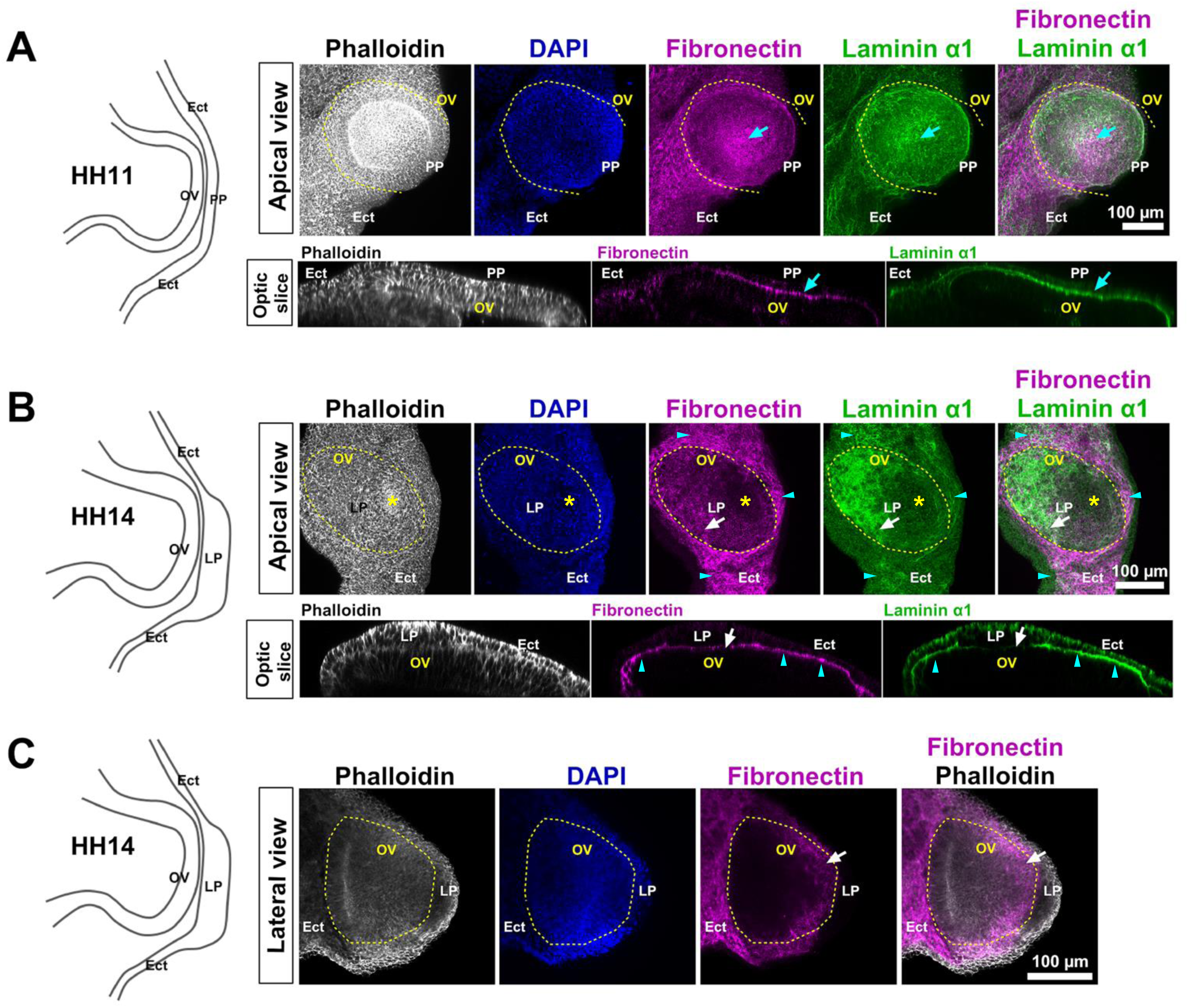
The ECM changes during lens placode formation. (A-C) On the left side, the line drawing represents the morphology of the lens placode at the stage of 3D-reconstructed fluorescence images and transversal optic slices. The optic tissue of HH11 to HH14 stage chick embryos were stained for actin filaments (phalloidin, white), Fibronectin (magenta) and Lamininα1 (green). Top row images are a view from the apical surface and the optical slices in the bottom fow are an orthogonal view of the YZ axis. (A) HH11 embryos display an intense labelling of Fibronectin and Lamininα1 between the optic vesicle and the pre-placodal ectoderm (cyan arrow). (B) At stage HH14 the lens placode is thickened, and actin accumulates in the apical surface where apical constriction begins (yellow asterisk). Fibronectin and Lamininα1 immunostaining is diffuse and punctate below the center of the placode (white arrows), contrasting with fibrillary organization in the non-placodal region (cyan arrowheads). The optical slice confirms that staining for both proteins between the lens placode (LP) and the optic vesicle (white arrow) is weaker under the placodal tissue. (C) Lateral view of the eye at stage HH14. Fibronectin immunostaining pattern is diffuse and punctate in the optic ECM (white arrow). Ect: non-placodal ectoderm; PP: pre-placodal ectoderm; OV: optic vesicle; LP: lens placode. The yellow dotted line delineates the optic vesicle (OV) at each developmental phase.

### Formation of optic ECM depend on BMP signaling in the lens placode

The dynamics of the optic ECM formation is strongly associated with the morphological changes at the onset of placodal differentiation. To verify if inhibition of placodal differentiation would also abolish the formation of optic ECM, we overexpressed a truncated form of the BMP receptor type 1 (tBMPr) separately in the lens placode and optic vesicle. This mutation eliminates the intracellular kinase domain and reduces the phosphorylation of Smad1/5/8 transcription factors even in the presence of BMP (Suzuki et al., 1994). With this, exogenous tBMPr acts as a dominant negative and inhibits the BMP signaling pathway in a cell-autonomous manner.

tBMPr overexpression at the pre-placodal ectoderm inhibited lens placode thickening and its invagination (Figure 2A, right column). At stage HH15, the electroporated pre-placodal ectoderm remained cuboidal while the lens placode formed normally in the control eye (Figure 2A, right column). In contrast, inhibition of BMP signalling in the optic vesicle had no detectable effect on the placode development and a mild effect on the optic cup (Figure 2A, left column). Despite normal placode formation, we observed a reduction in eye size in several embryos (data not shown). The formation of a smaller eye confirms that BMP signaling is necessary for the development of the optic vesicle (Rajagopal et al, 2009; Huang et al. 2015). These results also suggest that BMP signaling is required in the pre-placodal ectoderm for lens placode formation, but not for the optic vesicle morphogenesis.

**Figure 2.**
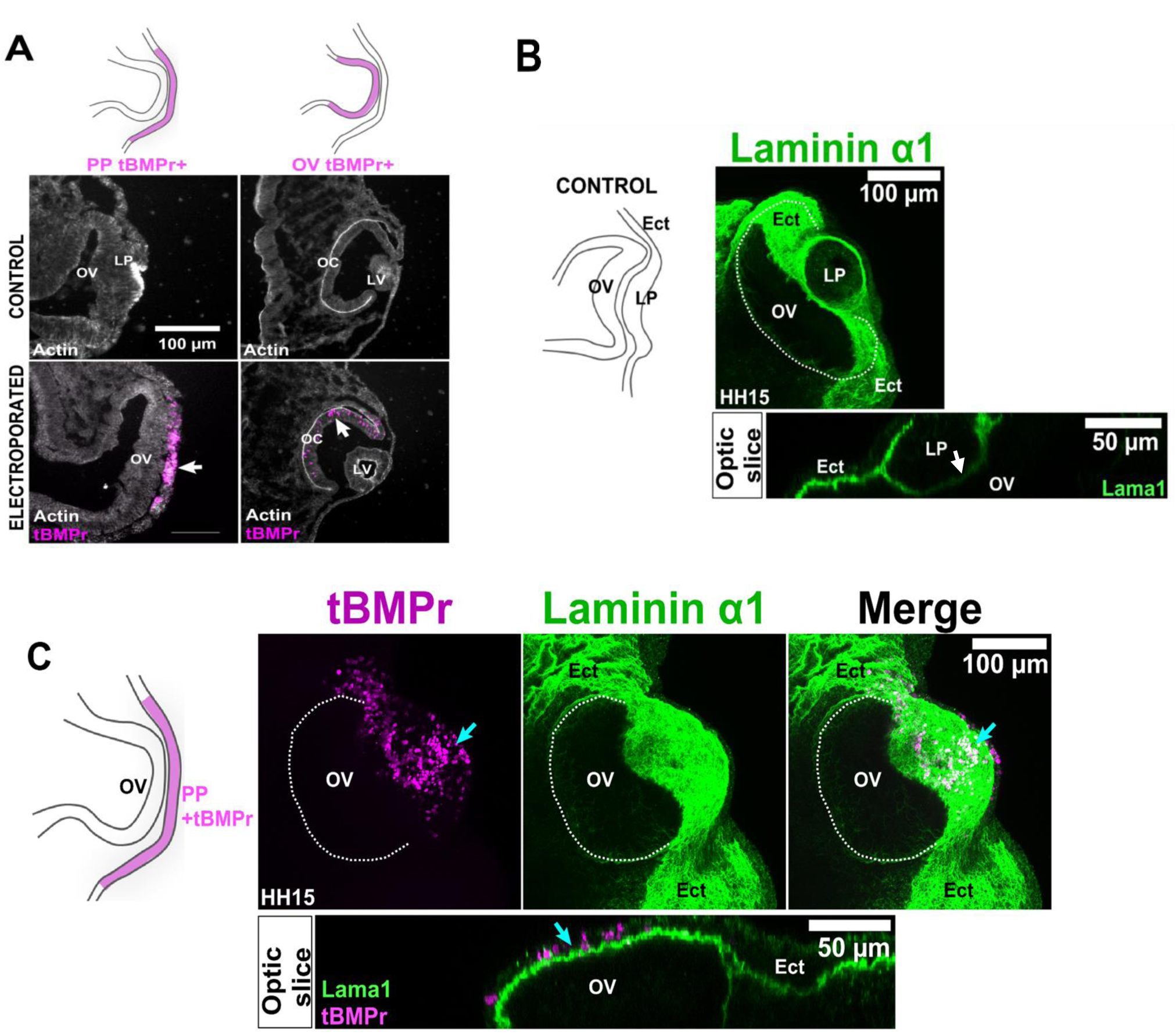
BMP signaling in pre-placodal cells I is required for ECM evolution. (A) Overexpression of truncated BMP receptor (tBMPr) in pre-placodal ectoderm (left) and optic vesicle (right) at HH9. The electroporated cells are seen in magenta (white arrow). Sections stained for actin with Phalloidin show that tBMPr in the pre-placodal ectoderm inhibits lens thickening, while its electroporation in optic vesicle (OV) does not affect lens placode formation (LV). (B-C) 3D images of chick embryo optic region at stage HH15 stained for Lamininα1 (green). Lamininα1 immunostaining pattern is fibrillar with intense labelling outside the optic region, between non-placodal ectoderm (Ect) and optic vesicle (white dotted line). YZ orthogonal slice shows weak Lamininα1 staining between the optic vesicle and the lens placode (LP, white arrow). (C) After tBMPr overexpression (magenta, cyan arrow) in the pre-placodal ectoderm, lens placode formation was inhibited. Lamininα1 immunostaining pattern is fibrillar with intense labelling both outside and inside the optic region. YZ orthogonal slice shows a side view of Lamininα1 staining between the optic vesicle (OV) and the tBMPr-positive ectoderm (cyan arrow). Ect: non-placodal ectoderm; PP: pre-placodal ectoderm; OV: optic vesicle; LP: lens placode; OC: optic cup; LV: lens vesicle; tBMPr: truncated BMP receptor.

Inhibition of BMP signaling in the pre-placodal ectoderm disrupted the above-described changes in Lama1 staining pattern (Figure 2C). Instead of differences between the placodal area and the rest of the cephalic region, we found that Lama1 labeling pattern remained fibrillar between the tBMPr-positive ectoderm and the optic vesicle (Figure 2C). These results suggest that the inhibition of BMP signaling in the pre-placodal ectoderm disrupts the mechanisms that change laminin distribution within the optic region.

On the other hand, when we overexpressed tBMPr in the optic vesicle, eye development was not affected and both Fn and Lama1 staining patterns are similar compared to the control. In other words, the staining was diffuse and punctate in the optic region (Figure 3A). In the non-placodal regions of the ectoderm, the ECM remained fibrillar (Figure 3A). The orthogonal slices show a more intense labeling of both Fn and Lama1 outside of the optic region compared to the optic region (Figure 3B). This stronger signal in the non-placodal region at the optic slice corresponds to the fibrillary pattern of both proteins in the apical view (Figure 3B, cyan arrowhead). Together, these results suggest that the distinct pattern of Lama1 and Fn in the optic region depends on BMP signaling in the placode but not in the optic vesicle. Furthermore, this suggests that optic ECM changes are modulated by lens placode cells.

**Figure 3.**
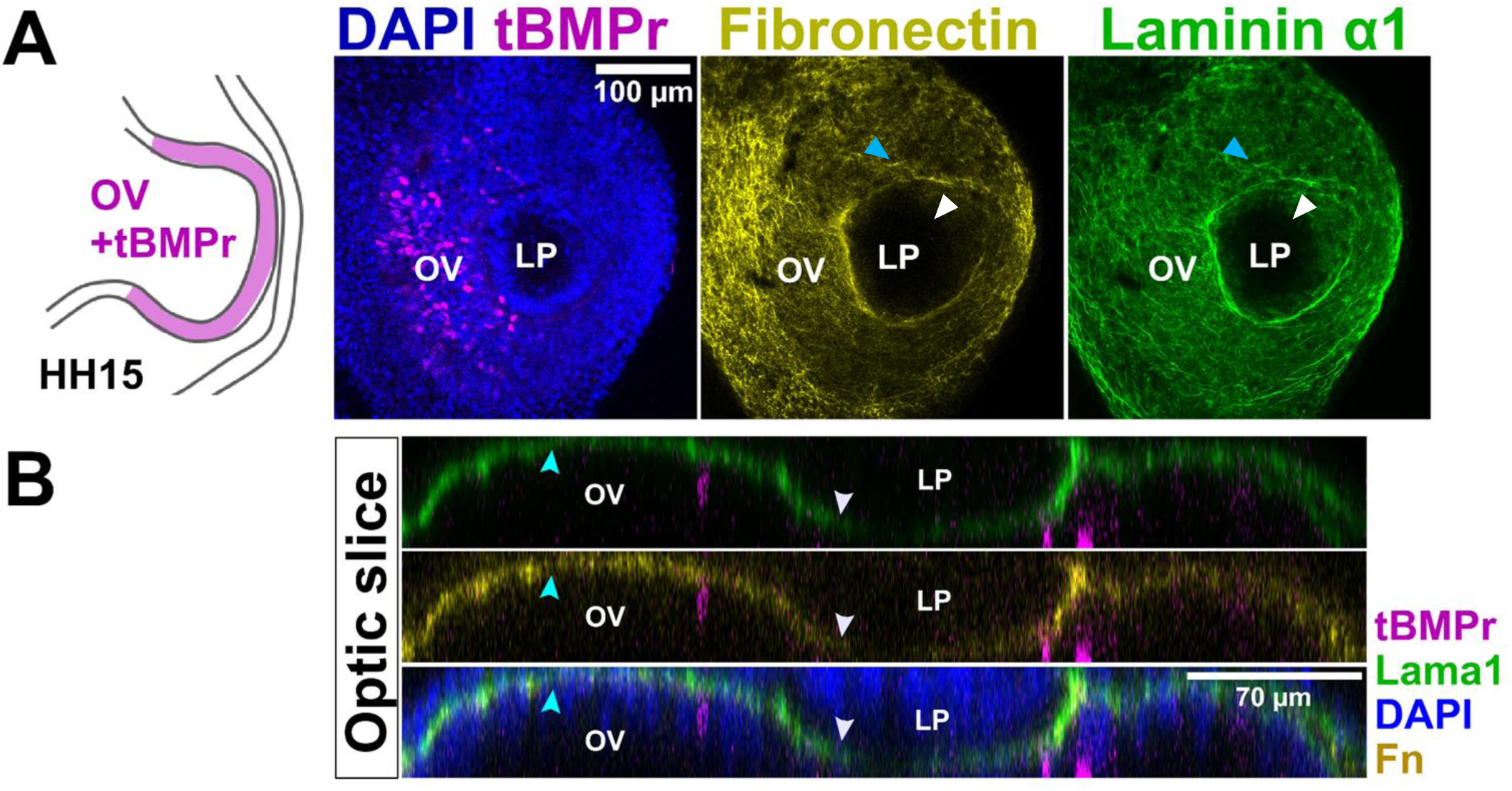
Inhibition of BMP pathway in the optic vesicle did not affect the evolution in Fibronectin and Lamininα1 staining pattern. (A) 3D fluorescence images of HH15 chick embryo optic region stained for Lamininα1 (green) and Fibronectin (yellow). After tBMPr overexpression in the optic vesicle (OV, magenta), the lens placode (LP) formed normally (white arrowhead). Both Laminin α1 and Fibronectin show a diffuse and punctate pattern between the lens vesicle and optic vesicle (white arrowhead). Outside the lens placode region, Fibronectin and Laminin staining is intense and fibrillar (cyan arrowhead). (B) YZ orthogonal slices show Lamininα1 and Fibronectin staining in non-placodal ectoderm (cyan arrowheads) and less intense labeling between electroporated optic vesicle and lens placode (white arrowheads). OV: optic vesicle; LP: lens placode.

### Lens placode modulates ECM within the optic region

To identify the main players that modulate the composition of optic ECM and investigate how the lens placode regulates it, we employed an unbiased approach using comprehensive transcriptomic data from mouse embryos. To identify which ECM genes are differently expressed in the lens placode, we analyzed multiple mouse embryo single-cell RNA sequencing (scRNA-seq) data. We first used the scRNA-seq dataset at Mouse Organogenesis Cell Atlas (Cao et al., 2019). These data were obtained from ∼2 million cells collected from 61 mouse embryos between stages E9.5–13.5 (Cao et al., 2019). Given the primary purpose of our study, we only used data from mouse embryos at stages E9.5 and E10.5 (total of 370,374 cells). We first filtered for *Pax6*-positive cells, which is expressed in both stages throughout the eye region of the mouse embryo (Cvekl and Zhang, 2017; Rowan et al., 2010). The resulting dataset of *Pax6*-positive cells at stages E9.5 and E10.5 generated 22 clusters. Of these, we identified 2 clusters with high expression of optic primordium markers such as DNA-binding transcription factors *Mitf*, *Rax*, *Lhx2*, *Prox1* and transient receptor potential channel *Trpm3*. This subgroup of optic cells has 555 cells. Upon regrouping, this subset formed 6 clusters (Figure 4A). To identify further the cell type represented on the 6 clusters, we analyzed the expression of optic vesicle and lens placode markers. We identified two clusters that have high levels of expression of early optic vesicle markers (cluster OV I and II: *Lhx2, Pax2, Rax*), two clusters with high retinal pigment epithelium markers expression (clusters RPE I and II: *Otx2* and *Mitf*), one cluster with high neural retina markers expression (clusters NR: *Vsx2*) and one cluster with high lens placode markers expression (cluster LP: *Cdh1*, *Prox1*, *Mab21l1*, *Maf*, and *Sfrp2*) (Figure 4D, Sup. Figure 3A) (Kakrana et al., 2017). The early optic vesicle clusters are formed mostly by stage 9.5 cells, and the other clusters are mostly composed by cells from stage E10.5 (Figure 4B).

**Figure 4.**
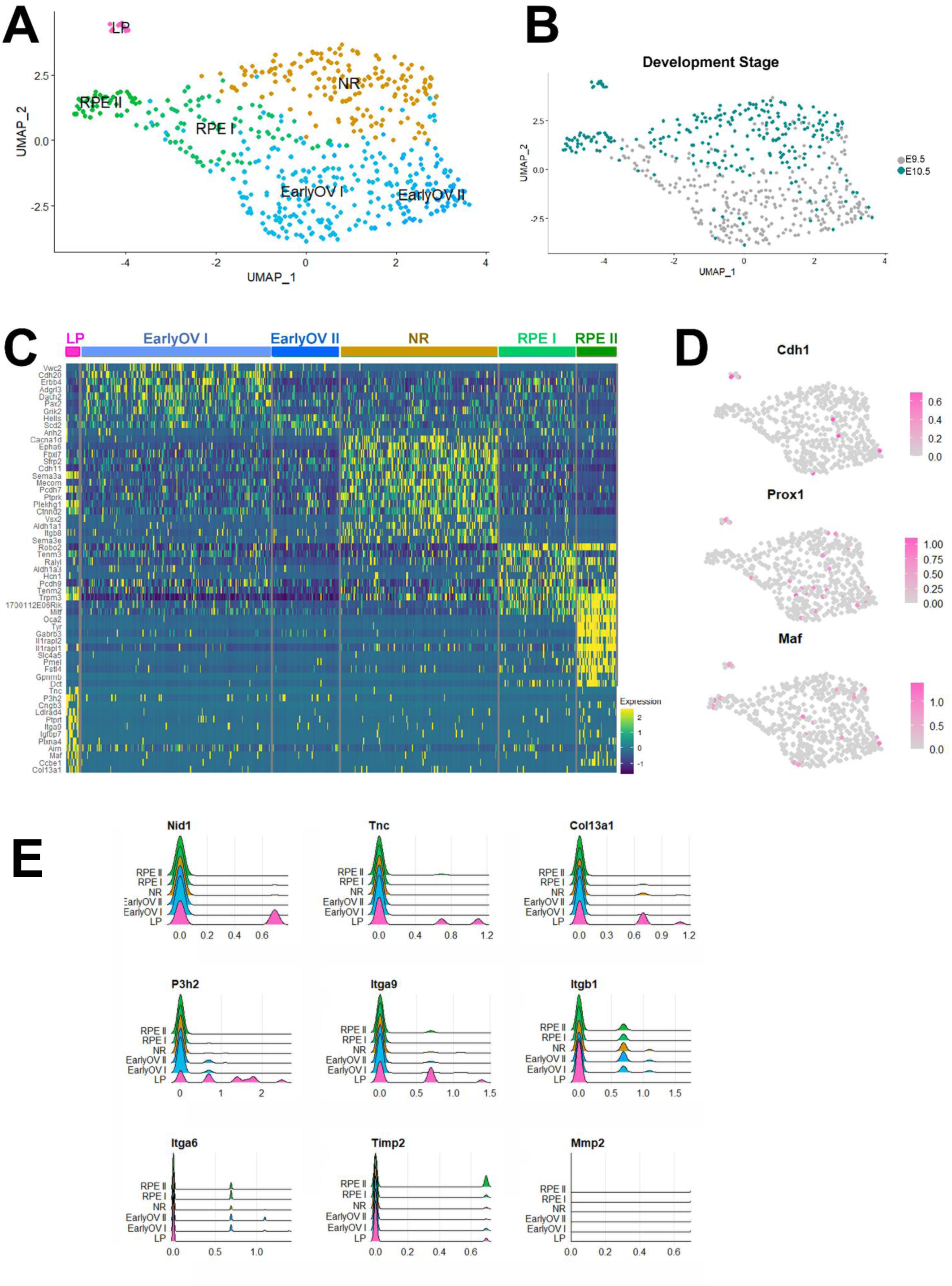
Mouse embryos lens placode cells express ECM-associated genes. (A) UMAP graph showing the clusters obtained after processing mouse embryo scRNAseq data. (B) Developmental stages mapped onto the UMAP graph. E9.5 cells are labelled grey, while blue represents E10.5 cells. (C) Heatmap of the top 10 differentially expressed genes in each cluster. High expression of optic vesicle markers, such as *Pax2, Vsx2,* and *Mitf*, classifies the clusters as early optic vesicle (OV), neuro retina (NR), and retinal pigment epithelium (RPE), respectively. The lens placode cluster (LP) was identified by the absence of optic vesicle markers expression and the high expression of lens placode markers. (D) The lens placode cluster exhibited high expression of *Cdh1*, *Prox1*, and *Maf*, while the optic vesicle clusters show low expression of these genes. (E) ECM-associated genes expression levels in each cluster. The y-axis represents the number of cells with a specific level of expression (x-axis). *Nid1, Tnc, Col13a1, P3h2*, and *Itga9* exhibit higher expression in lens placode cluster cells than in the other clusters (LP, pink). *Itgb1* and *Itga6* are expressed by some optic clusters (RPEI/II, EarlyOVI/II, NR) but are absent in the lens placode cluster. *Mmp2* expression is absent in all cell types.

The lens placodal cluster has only stage E10.5 cells and would morphologically correspond to invaginating placode/lens pit (Figure 4B). The 20 most differentially expressed genes (DEGs) for each cluster are listed in Sup. Figure 3B. Among the DEGs of the lens placode cluster, we found several genes associated with the extracellular matrix, such as *Nidogen1* (*Nid1*, also known as *Entactin*), *Tenascin-C* (*Tnc*), *Leprecan-like protein 1* (*P3h2* or *Leprel-1*), and *Collagen Type XIII Alpha 1 Chain* (*Col13a1*) (Figure 4E). Of these, Nid1 was the most differently expressed ECM gene.

Several DEGs found in lens placode cluster were also identified as down-regulated in microarray data from mouse embryos harboring a lens-specific depletion of *Pax6* (Huang et al., 2011). These embryos do not develop lenses and the *Pax6*-knockout cephalic epithelia does not express lens development regulatory genes such as *Prox1* and *Maf*. In this paradigm, *Tnc*, *P3h2*, *Col13a1* were also reported as downregulated (Huang et al., 2011). These results strengthen our conclusions from scRNA-seq analysis and suggests that the expression of *Tnc*, *P3h2* and *Col13a1* are ECM-related genes required for lens placode differentiation.

We then proceeded to validate some of the genes identified in the above scRNA-seq analysis. We focused on Nid1 and Tnc. In E9.0 mice, Nid1 was detected in the basal lamina of all neural and non-neural ectoderm prior to lens placode thickening (Figure 5A). However, contrary to what we saw with Laminin and Fibronectin, there was no difference in Nid1 labeling pattern or intensity in the placodal region. Nid1 is present in the ECM between placodal ectoderm and optic vesicle and in the non-placodal ectoderm. The intensity and pattern of Nid labelling in the optic region was similar to the non-optic regions of the cephalic ectoderm.

**Figure 5.**
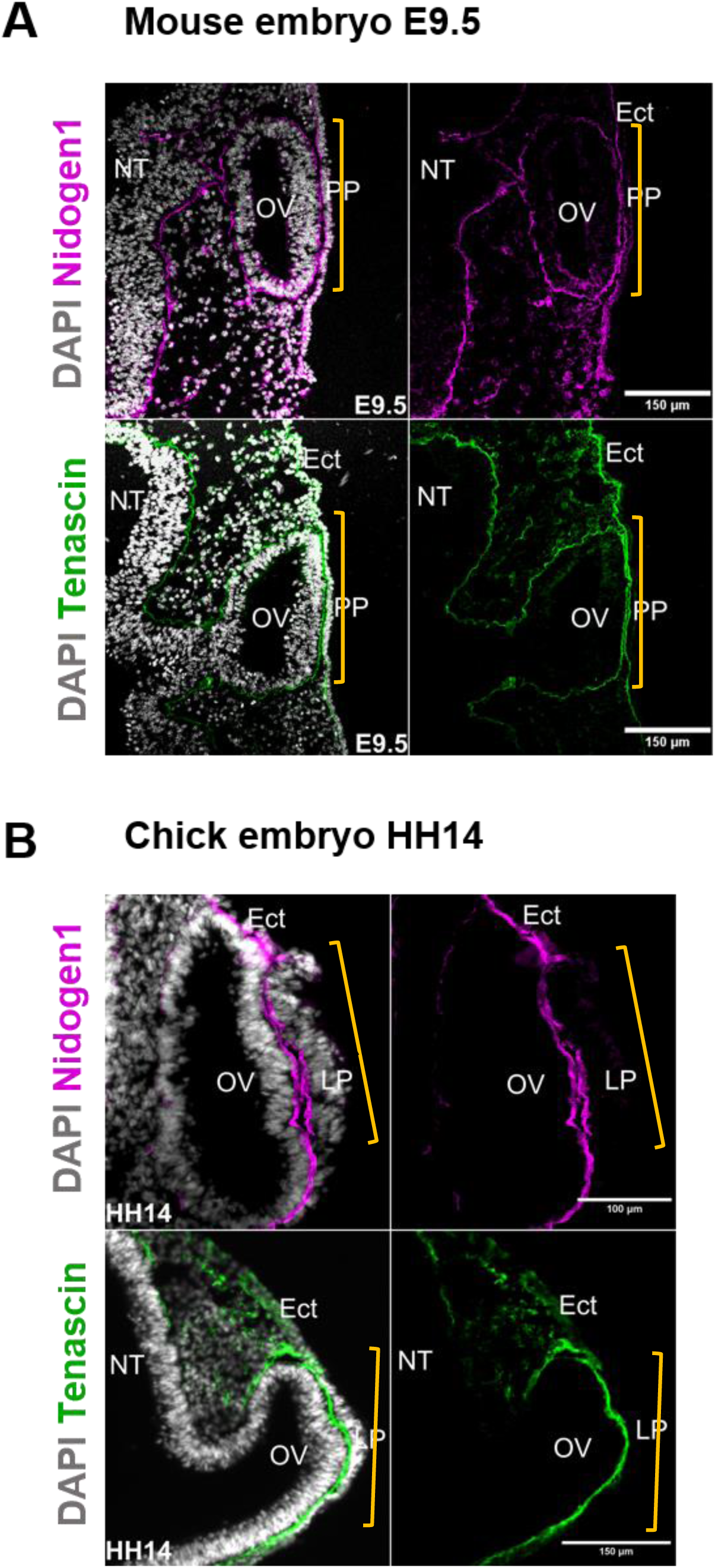
Tenascin and Nidogen1 are in the ECM between lens placode and optic vesicle. (A) Nidogen1 (Nid1, magenta) and Tenascin (Tnc, green) immunostaining in E9.5 mouse embryo sections. Nid1 and Tnc labeling is intense between the pre-placodal ectoderm (PP) and the optic vesicle (OV). The yellow brackets delimit the optical region. (B) Nidogen1 (Nid1, magenta) and Tenascin (Tnc, green) immunostaining in HH14 chick embryo. Both proteins are detected in the basal domain of the lens placode (LP) and the non-neural ectoderm (Ect). The yellow brackets delimit the optical region. Ect: non-placodal ectoderm; NT: neural tube; OV: optic vesicle; LP: lens placode; PP: pre-placodal ectoderm.

Similarly, Tnc is also present in the basal ECM underlying all the epithelia: the neural tube, optic vesicle, pre-placodal ectoderm, and non-placodal ectoderm. In the chick embryo, Tnc and Nid1 were also in the optic ECM (Figure 5B). Notably, Tnc labeling around the optic vesicle and in the basal region of the lens placode is more intense compared to other regions of the cephalic ectoderm (Figure 5B). Together, these results confirm that Nid1 and Tnc are present in the optic ECM in both chick and mouse embryo. In addition, the scRNAseq data suggests that the main source of both molecules is likely the lens placode.

To investigate if the same ECM-associated genes were also expressed earlier, we also analyzed scRNA-seq data from the dissected E9.5 eye region (Yamada et al., 2021) (Sup. Figure 4). This analysis also suggests that *Tnc*, *P3h2*, *Col13a1* are highly expressed in lens placode cluster compared to optic vesicle clusters (Sup. Figure 4D). Thus, analysis of both scRNA-seq datasets suggests that the lens placode expresses proteins that contributes to the embryonic optic ECM.

### Lens placode inhibits gelatinase activity in the optic region

Our *in silico* analysis further suggested that *Timp2*, an MMP2 inhibitor, is expressed in optic vesicle and lens placode clusters (Figure 4E). Furthermore, MMPs are not significantly expressed in the optic tissue during lens placode formation (Figure 4E).

To confirm that *Timp2* is indeed present and whether its expression is tissue and stage-specific, we performed an *in situ* hybridization assay before and during chick lens placode formation (Figure 6A). In the pre-placodal stage (HH11), *Timp2* is expressed within the distal region of the optic vesicle. We did not detect *Timp2* expression in the pre-placodal ectoderm and other regions of the head surface ectoderm. In the placode stage (HH14), *Timp2* is expressed throughout the placode during its thickening. In contrast, the surrounding surface ectodermal cells do not express *Timp2*. Thus, *Timp2* increases with the onset of placodal development specifically in the placode, suggesting that its activity is closely associated with placodal morphogenesis. This increase could downregulate metalloprotease activity in the optic region.

**Figure 6.**
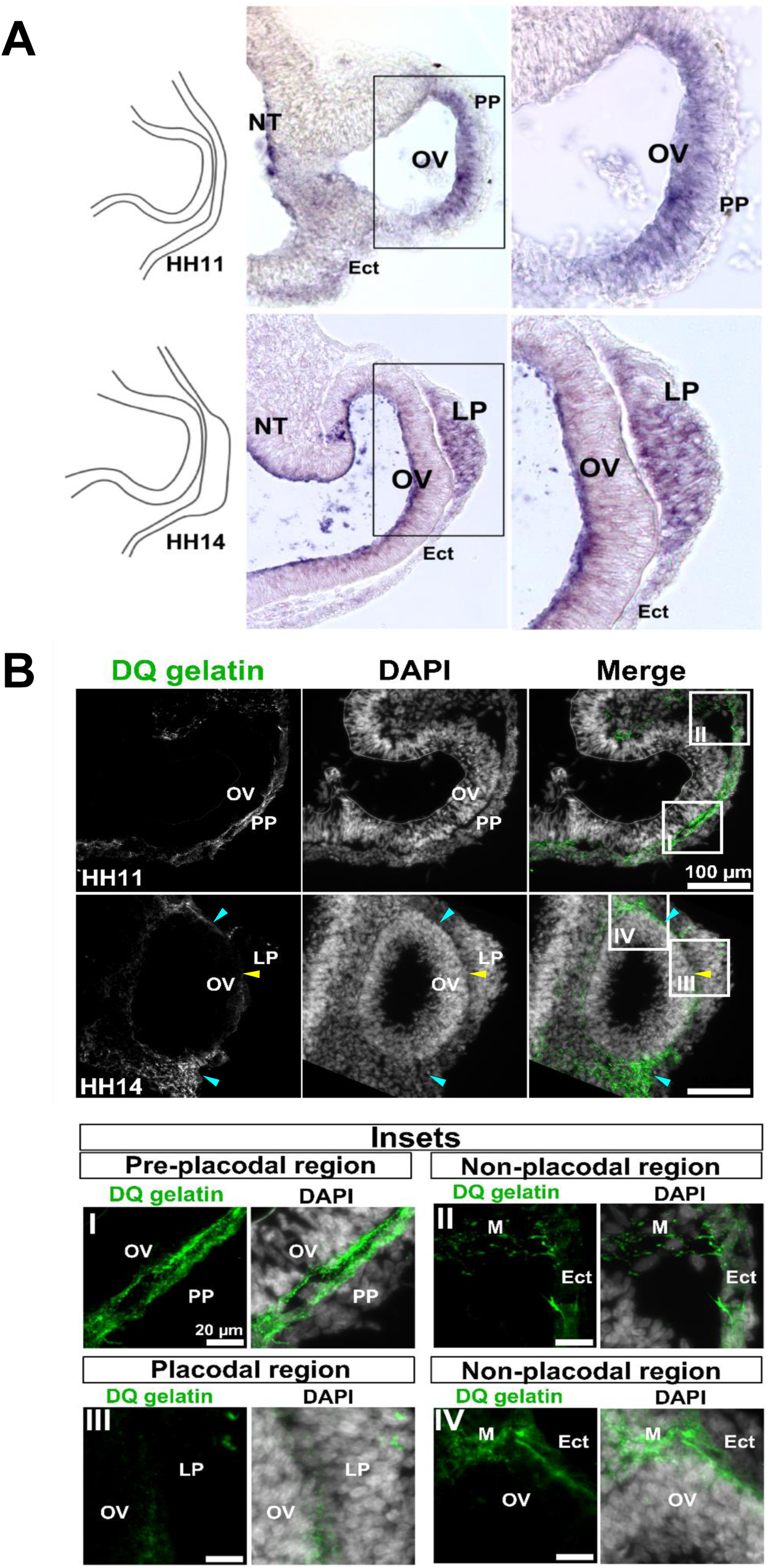
*Timp2* expression in the lens placode increases, while gelatinase activity decreases during lens placode formation. (A) Sections of chick embryo at stage HH11 and HH14 (schematic representations on the left) with *Timp2* expression labelling by *in situ* hybridization. At stage HH11, *Timp2* is expressed in the optic vesicle (OV). At stage HH14, *Timp2* is expressed in lens placode cells (LP) and is absent in other tissues. (B) *In situ* zymography on frozen sections of chick embryos shows gelatinase activity through DQ-gelatin labelling. At stage HH11 (upper row), intense DQ-gelatin labelling is observed between the pre-placodal ectoderm and the optic vesicle (I). Gelatinase activity is also present in the non-placodal ectoderm (II, Ect). At stage HH14 (lower row), DQ-gelatin labelling is only present outside the optic region (IV, cyan arrowheads). There is minimal protease activity between the lens placode and the optic vesicle (III, yellow arrowhead). Insets corresponds to higher magnifications. Ect: non-placodal ectoderm; PP: pre-placodal ectoderm; OV: optic vesicle; LP: lens placode.

To test this hypothesis, we followed the dynamics of metalloprotease activity during placodal growth. We performed *in situ* zymography, in which a quenched fluorescently-labeled gelatin is digested by active gelatinases, such as MMP2 and MMP9 (Gkantidis et al., 2012; Kalev-Altman et al., 2020; Mook et al., 2003; Porto et al., 2009). After digestion, the fluorescence signal appears and labels the sites with gelatinase activity. At early stages (HH11, before lens placode thickening), we observed a high gelatinase activity between pre-placodal ectoderm and the optic vesicle (Figure 6B). Higher magnification of the optic region shows two DQ-gelatin labelled lines, near the apical surface of the optic vesicle and the basal surface of pre-placodal ectoderm. Gelatinase activity was also present in the cephalic ectoderm around optic region (Figure 6B). At later stages (HH14, after lens placode thickening), gelatinase activity decreased significantly between the lens placode and the optic vesicle (Figure 6B). In contrast, it remains throughout the basal region of the non-optic surface epithelia at all stages (Figure 6B). This result demonstrates that gelatinase activity is excluded from the lens placode during its thickening. Notably, gelatinase activity is complementary to *Timp2* expression sites.

### Modulation of metalloprotease activity in optic ECM depends on BMP signaling

Since Fn and Lama1 organization changes depend on BMP signaling, our next question was whether *Timp2* expression and gelatinase activity also require BMP signaling in the pre-placodal ectoderm. Thus, we analyzed *Timp2* expression at stage HH14 after tBMPr overexpression at the pre-placodal ectoderm (Figure 7A). Inhibition of BMP signaling in the pre-placodal tissue arrested lens placode formation and reduced *Timp2* expression (Figure 7A). In contrast, the control eye developed normally, displaying expression of *Timp2* specifically at placodal cells (Figure 7B)..

**Figure 7.**
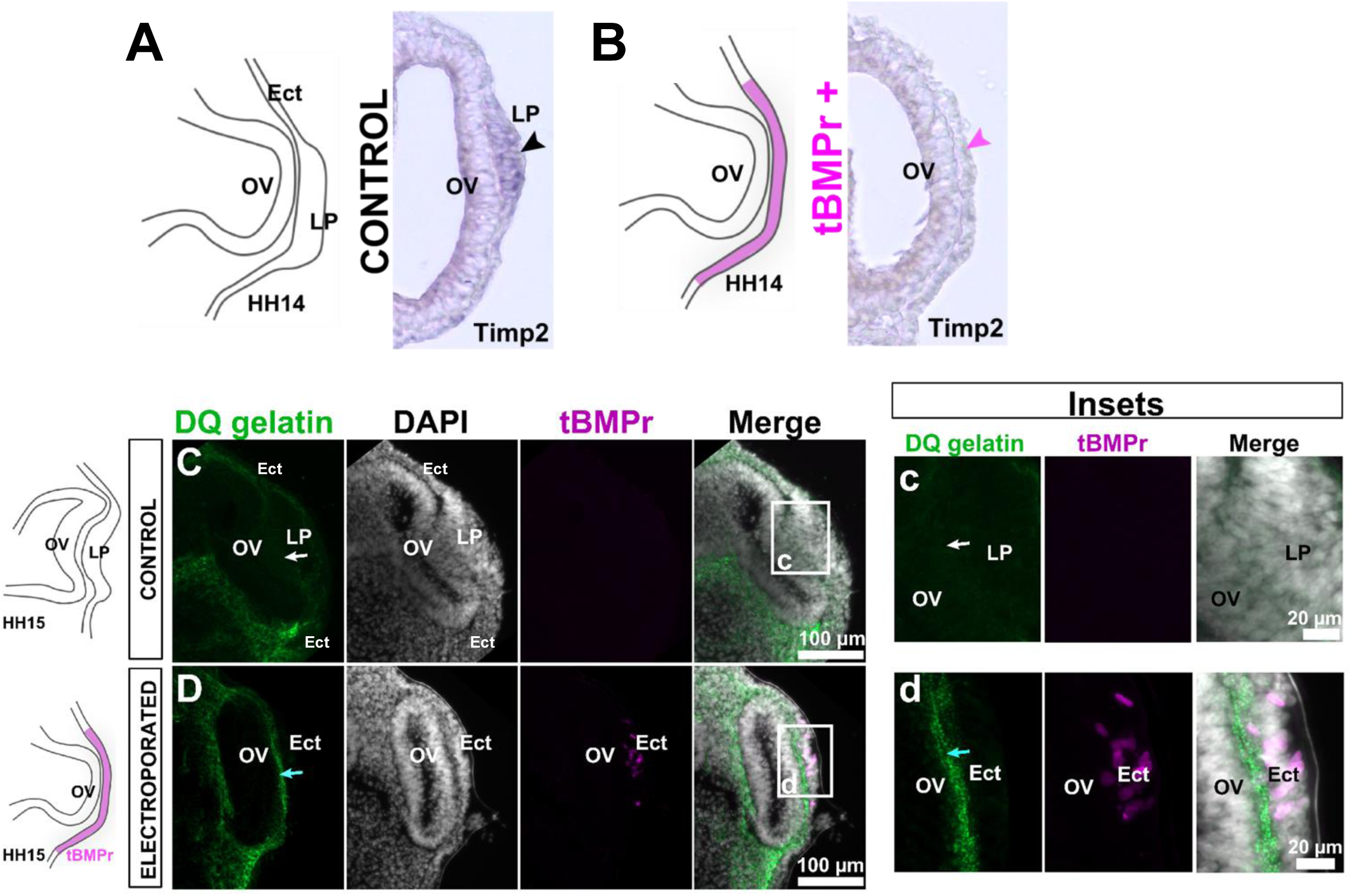
BMP signaling in the pre-placodal ectoderm is required *Timp2* expression and reduction of gelatinase activity. (A-B) I*n situ* hybridization for *Timp2* in stage HH14 chick embryos. (A) In the control eye *Timp2* is expressed specifically in the lens placode (black arrowhead). (B) When tBMPr was overexpressed in the pre-placodal ectoderm, the lens placode failed to thicken and there is no expression of *Timp2* (pink arrowhead) (C-D) *In situ* zymography assays. (C) The control eye at stage HH15 shows no gelatinase activity in the optic ECM (white arrow), between lens placode (LP) and optic vesicle (OV). (c) Higher magnification of lens placode shows the absence of DQ-gelatin labelling in the optic region (white arrow). (D) In the electroporated eye, lens placode formation was inhibited and gelatinase activity was detected between the optic vesicle (OV) and electroporated ectoderm (Ect) (cyan arrow–(d) Higher magnification of the electroporated eye shows an intense DQ-gelatin labelling in the optic ECM.

Next, we performed *in situ* zymography after tBMPr overexpression in the pre-placodal tissue (Figure 7D). Inhibition of BMP signaling arrested lens development. Importantly, we observed an intense labeling of DQ-gelatin between the tBMPr-positive ectoderm and the optic vesicle, indicating that reduction of ECM protease activity requires placodal development (Figure 7D). Together, these results show that *Timp2* expression and inhibition of gelatinase activity during lens placode thickening both depend on BMP signaling in the pre-placodal ectoderm.

## DISCUSSION

Here, we analyzed for the first time 3D-dynamic changes of multiple novel as well as established optic ECM proteins during the critical time window of the early vertebrate eye development. Our main findings are represented as a graphical abstract in Figure 8. Lama1 and Fn change their staining pattern when the lens pre-placode ectoderm undergoes conversion into the placodal morphology (Figure 8A and B). Previous studies also describe changes in the ECM-staining during lens placode formation. In both chick and mouse embryos, PAS-staining (Periodic acid-Schiff staining detects polysaccharides such as glycogen, and mucosubstances such as glycoproteins) between the placodal ectoderm and the optic vesicle becomes more intense during placode formation (Hendrix and Zwaan, 1975; Huang et al., 2011). Furthermore, after placode thickening, highly acidic glycosaminoglycans concentrate specifically at the optic vesicle apical surface (Hilfer et al., 1981). Together, these data show that during lens placode development there is a temporally and spatially regulated change in the ocular ECM. Our results confirm that these changes occur only in the optic ECM and not at the surrounding ECM that underlies other portions of the surface ectoderm. Notably, both Fn and Lama1 staining patterns remain fibrillary outside the optic field.

**Figure 8.**
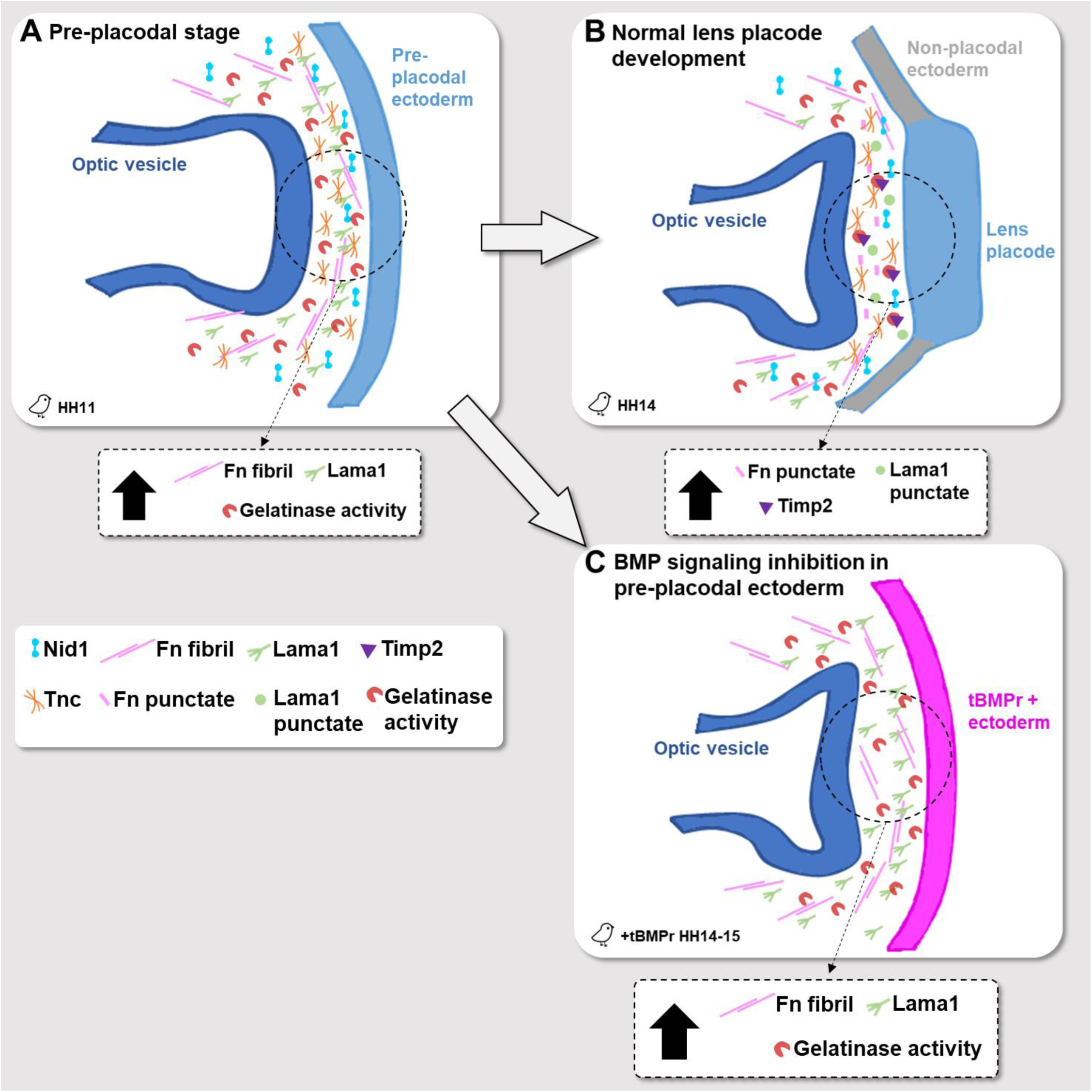
Working model for ECM changes in the optic region. At the pre-placodal stage (HH11 top left), Fibronectin and Lamininα1 display a fibrillar pattern in the optic region and in the surrounding non-placodal ectoderm. Nidogen1, Tenascin and gelatinase activity are also present and evenly distributed throughout the cephalic ectoderm. At later stages, when the lens placode thickens (HH14, top right), Fibronectin and Lamininα1 staining pattern become more diffuse and punctate between the optic vesicle (dark blue) and the thickened lens placode (light blue). In contrast, under the non-placodal ectoderm (grey) the pattern remains fibrillar for both proteins. While Nidogen1 remains similar inside and outside the optic region, Tenascin is stronger in the optic ECM. Further, the lens placode expresses the gelatinase inhibitor *Timp2*. Gelatinase remains active only in the non-placodal ectoderm, outside the optic region. Without BMP signaling (lower right), the lens placode does not develop, and the ECM is not remodeled. With the inhibition of BMP in the pre-placodal ectoderm (pink), Fibronectin and Lamininα1 maintain the fibrillary pattern, *Timp2* is not transcribed, and gelatinase activity remains high throughout the ectoderm.

We identified, for the first time, the presence of Tnc in the early optic ECM at both the pre-placodal and placodal stages. Tnc is a glycoprotein that modulates adhesion through cell-matrix interactions (Midwood et al., 2016). It binds to Fibronectin and reduces the interaction between Fibronectin and Syndecan-4. This inhibits cell-matrix adhesion, as it disturbs formation of focal-adhesions and stress fibers (Chiquet-Ehrismann and Tucker, 2011; Midwood and Schwarzbauer, 2002; Van Obberghen-Schilling et al., 2011). In the embryo, Tnc is highly expressed in regions of intense morphogenesis during organ formation (Chiquet-Ehrismann and Tucker, 2011; Yoshida and Aoki, 2014; Midwood et al., 2016). In the optic region, Tnc has been previously described around the lens capsule of the eye (DeDreu et al., 2021). It is also expressed in mouse retina at stages E13.5 to 18.5 and regulates retinogenesis (Besser et al., 2012). Finally, expression of different Tnc isoforms is regulated by Pax6 (von Holst et al., 2007).

Another ECM glycoprotein that is constantly associated with the optic matrix during placodal growth is Nid1 (also known as Entactin). Nidogens are encoded by one gene in invertebrates, two genes in mammals and chick (*Nid1* and *Nid2*), and four genes in zebrafish (Bryan et al., 2020; Zhang et al., 2022). Nid1 is expressed in mouse lens placode at stages E9.5, E10.5, and E.12 (Dong and Chung, 1991; Grindley et al., 1995). Accordingly, our data shows that Nid1 is also present in the chick eye at placodal stages. Nid1 and Nid2 have important role in basement membrane assembly and homeostasis in optic cup morphogenesis, limb, heart and lung development (Bader et al., 2005; Böse et al., 2006, Bryan et al., 2020). Single Nid1 and Nid2 mutants do not show severe phenotypes (Zhang et al., 2022). Mice mutant for Nid1 do not show obvious alterations during embryogenesis and embryonic basement membrane formation (Böse et al., 2006; Murshed et al., 2000; Zhou et al., 2022). Although the basement membrane has important mechanical roles and regulates tissue shape, Nid1 itself has not been associated to intense morphogenesis process. Given the different roles of Tnc and Nid1 in the cell-matrix relationship, and the difference in optic and non-optic staining pattern shown here, we propose that Tnc might play a more relevant role in the lens placode differentiation and shape definition compared to Nid1.

Importantly, our data suggests that changes in optic Lama1 and Fn depend on lens placode formation. Inhibition of BMP signaling in the placode, but not in the optic vesicle, arrested ECM changes, indicating that these changes are driven by the lens placode (Figure 8C). Indeed, the transcriptomic profile of the optic tissues indicates that the placode contributes with unique components of the ECM. Further, lens-specific *Pax6* knockout mice fail to form lens placode, optic ECM deposition decreases, and several ECM-associated genes are downregulated (Huang et al., 2011). Interestingly, some of these downregulated genes (e.g., *Tnc*, *P3h2* and *Col13a1*) were also identified by our transcriptomic analysis to be characteristics of lens placode clusters. This data strengthens our hypothesis that lens placode differentiation is necessary for modulation of optic ECM.

Changes in optic ECM have been shown to be important for morphogenesis of the optic vesicle. Invagination of the optic vesicle starts at HH13 and converts it into the double-layered optic cup. During HH11-HH13, highly acidic glycosaminoglycans accumulate at the apical cell surfaces of the optic vesicle. Pharmacological inhibition of glycoconjugate synthesis inhibits optic vesicle invagination. In contrast, lens placode development is not interrupted, and can even form a small lens vesicle (Hilfer et al., 1981). This data shows that interference with changes in the optic ECM prevents invagination of the optic vesicle but does not interfere with the formation of the lens placode. The importance of the ECM in epithelial morphogenesis has been well characterized in the more posterior otic placodes as well. Perturbation of otic ECM with Laminin and Integrins antibodies interrupted otic placode invagination (Visconti and Hilfer, 2002).

Optic vesicle invagination also requires the presence of the pre-placodal ectoderm but not of the mature placode. This requirement is restricted to a specific window of time. Surgical removal of the chick pre-placodal ectoderm at HH11, abolishes optic vesicle morphogenesis. In contrast, when the lens placode is removed after HH13, the optic cup forms normally, suggesting that optic vesicle morphogenesis no longer requires the presence of the lens placode (Hyer et al., 2003; Oltean et al., 2016). However, the combination of placodal removal with collagenase treatment at HH13 also inhibits the invagination of the optic vesicle, indicating that the ECM under the HH13 placode is fundamental for optic vesicle invagination (Oltean et al., 2016). In view of our present data, we interpret that between HH11-13, the placode is the main tissue that alters the ECM to support optic vesicle morphogenesis. We propose that, while the placode grows, it defines the biomechanical properties and composition of the optic ECM that are essential for formation of the optic cup. Once the optic ECM acquires a specific composition and mechanical characteristics that are essential for the formation of the optic cup, the placode itself is not required for morphogenesis of the optic cup.

The full spectrum of molecular factors responsible for optic ECM remodeling remain unknown. The molecular mechanisms that act in ECM modulation during early eye development can be multifactorial. One of the main mechanisms of ECM remodeling is ECM degradation through matrix metalloproteinases (MMPs) activity (Diaz-de-la-Loza et al., 2018; Winkler et al., 2020). MMPs cleave ECM proteins and change matrix organization, release matrix-bound growth factors and other ECM fragments. The role of MMPs in morphogenesis has been well described in various organisms. In *Drosophila*, for example, the elongation of wings and legs involves a columnar-to-cuboidal cell shape change. This cell height reduction depends on ECM degradation through MMP1/2 activity. Inhibition of MMP1/2 activity maintains the columnar cell shape and disrupt elongation of wings and legs (Diaz-de-la-Loza et al., 2018).

Here we show that*Timp2*, an MMP2 inhibitor, is highly and specifically expressed in the lens placode during its thickening. Further, our *in silico* analyses show that MMP2 expression is low in optic tissues in early eye development. Finally, gelatinase activity decreases specifically in the optic ECM during placode thickening (Figure 8A, B). Together, these data suggest that MMP2 activity might be tightly regulated in space and time during lens placode formation. We propose that the increase in *Timp2* expression is probably an additional mechanism that inhibits MMP2 activity specifically in the optic ECM during placodal development.

A potential explanation for the importance of inhibiting the activity of metalloproteases such as MMP2 during early optic development would be to avoid premature entry of neural crest cells (NCC). NCC restrict development of lens fate outside pre-placodal region, and their absence results in ectopic lens formation (Bailey et al., 2006; Grocott et al., 2011). Previous studies suggested that the optic vesicle act as a physical barrier that prevents the migration of NCC in the pre-placodal region (Grocott et al., 2011). During pre-placodal stages (HH11-12), NCC migrate from the dorsal region of the neural tube and reach the lateral and proximal regions of the optic vesicle. They migrate around the optic region and towards the non-placodal ectoderm. This migratory pattern ensures that the space between the pre-placodal ectoderm and the distal portion of the optic vesicle remains unaffected by NCC presence during early lens development (Bailey et al., 2006; Grocott et al., 2011; Theveneau and Mayor, 2012). NCC only approaches the lens epithelia at later stages, after lens vesicle formation (Creuzet et al., 2005; Weigele and Bohnsack, 2020). Since the NCC require MMP2 and MMP9 activity to migrate (Kalev-Altman et al., 2020), it is likely that the inhibition of these gelatinases would further inhibit NCC migration. In addition to the gelatinase activity, NCC migration remodels fibronectin to a fibrillar-pattern. This process is called fibrillogenesis, and is critical for the neural crest cells migration (Martinson et al., 2023). In this context, the punctual pattern of fibronectin between the placode and the optic vesicle could also contribute to inhibit NCC entry to this region during early lens development.

Taken together, the present data show that the lens placode plays an active role in remodeling the optic ECM during early eye development. We highlight the crucial role of lens placode differentiation in orchestrating optical ECM characteristics and remodeling. Our research uncovered the expression of lens placode-specific ECM genes, including *Timp2*. *Timp2* temporal and spatial expression pattern correlated closely with reduction of gelatinase activity under the placodal region. This strongly suggests that Timp2 functions as an inhibitor of MMP2 within the optical ECM. The downregulation of Timp2 and upregulation of gelatinase activity in the absence of BMP signaling provide additional evidence that optical ECM modulation is intricately linked to placode formation. We thus propose that optic ECM remodeling depends on lens placode differentiation. In this scenario, evolution of the ECM could be relevant for maintenance of lens fate and morphogenesis of the optic cup.

## MATERIALS AND METHODS

### Chick and mice embryos

We obtained fertilized Leghorn chicken eggs from Granja Yamaguishi, at São Paulo state (for embryos used in experiments in the University of São Paulo) and Henry Stewart & Co., at Norfolk (for embryos used in experiments in the University of Oxford). The eggs were incubated at approximately 37.7 °C and 50% relative humidity for 34 to 50 hours to obtain embryos at different stages based on Hamburger and Hamilton, 1992. The wild-type mice embryos at stage E9.5 and E10.5 were obtained from pregnant CD-1 females from Charles River (Cvekl Laboratory, Albert Einstein College of Medicine). The experiments in chick embryos have been approved by the Ethics Committee at the Biomedical Sciences - USP (CEUA#9506131021) and the experiments in mouse embryos are approved by the Institute of Animal Studies at the Albert Einstein College of Medicine (# 00001533).

### *In ovo* chick embryo electroporation

Chick embryos were electroporated at stage HH9 in optic vesicle and pre-placodal ectoderm. The plasmid that was used was pCI-DN-BMPr1-H2B-RFP (donated by Marcos Simões-Costa, Cornell University at Ithaca, USA). The plasmid contains a truncated sequence of the BMP receptor type 1 from chicken with a truncation in the C-terminal portion. Prior to electroporation, the plasmid was diluted in H_2_O to a concentration of 1,5-2,5 μg/μl.

### Immunofluorescence of whole-mount embryos

Chick and mouse embryos were staged and dissected in PBS, fixed in paraformaldehyde 4% for 25-30 minutes, and washed 3 times for 10 minutes in PBS at room temperature. After fixation and washing, the embryos were permeabilized in 1% Triton X-100 in PBS for 30 minutes. We incubated in a block solution of 3-5% bovine serum albumin (BSA) in PBS for 3h at room temperature. The primary antibodies used were: mouse monoclonal anti-chicken Fibronectin (DSHB, clone B3/D6, diluted 5-1 ng/ml), rabbit polyclonal anti-Lamininα1 (Sigma, L9393, diluted 1:60), mouse anti-chicken Tnc (DSHB M1B4, diluted 1:50), mouse anti-chicken Nid1 (DSHB 1G12, diluted 1:50). All primary antibodies were diluted in PBS with 1% BSA and 0.1% Triton X-100 (Rifes and Thorsteinsdóttir, 2012). The secondary antibodies used were: anti-mouse IgG Alexa Fluor 568, anti-mouse IgG Alexa Fluor 647 and anti-rabbit IgG Alexa Fluor 488. We also stained the embryos with phalloidin coupled to different fluorophores (Thermo Life Scientific, 1:100) and DAPI (Invitrogen, 1 mg/ml, diluted 1:1000). Whole-mount embryos imaging were acquired on a Zeiss LSM-780 NLO (CEFAP, ICB – USP, FAPESP 2009/53994-8), on Zeiss 880 Airyscan fast (Srinivas Laboratory, University of Oxford) and on Leica SP8 Inverted DMi8 (Analytical Imaging Facility – Albert Einstein College of Medicine, SIG # 1S10OD023591-01 and Cancer Center P30CA013330). In Zeiss 880 Airyscan fast, we used a 40x/1.2 oil immersion lens (Objective C-Apochromat 40x/1.20 W Korr M27) with 1.5 digital zoom. In Leica SP8, we used a 20x 0.75 Air PLAPO and a 40x 1.3 NA oil immersion PLAPO objectives.

### Immunofluorescence of embryo cryosection

After fixation, embryos were cryoprotected in 20% sucrose, embedded in O.C.T compound (Tissue-Tek) with 20% Sucrose (1:1), and cryosectioned at 10-12 µm. We blocked the sections with 3-1% BSA for 1 hour and used the same set of primary antibodies described previously. The sections were mounted on glass slides using Vectashield mounting medium with DAPI. Images were taken using a Zeiss Widefield Axiobserver microscope, equipped with Zeiss digital camera Axiocam 208 and ApoTome.2. Images were acquired using the Zen blue software and, later, processed in Fiji software.

### *In situ* zymography using DQ-gelatin

For the analysis of the protease activity assay, we used DQ-gelatin (E12055, Molecular Probes, Eugene, OR) as a substrate in unfixed cryosections. The solution was prepared from the DQ-gelatin reagent 1mg/ml in water, diluted 1:10 in 50mM TrisCaCl_2_ (Porto et al., 2009). Before using, we heated the solution for 5 minutes at 37°C, quickly vortexed twice, and put it in an ultrasonic bath for 10 minutes. The slides with DQ-gelatin were incubated for 8 to 12 hours at 37°C in a humid chamber. We interrupted the reaction with PFA 4% and vigorously washed the slides 10 times with PBS. We mounted the sections using Vectashield mounting medium with DAPI (Electron Microscope Sciences #17989-20).

The DQ-gelatin is a quenched fluorogenic gelatin substrate. The fluorescent FITC molecule becomes exposed upon proteolytic digestion. It is mainly used to detect MMP2 (gelatinase B) and MMP9 (gelatinase A) activities, but other MMPs with weaker gelatinolytic activity, serine, cysteine or aspartic proteinases might contribute to the signal (Snoek-van Beurden et al., 2005). To verify if our zymography labeling of the chick embryo optic region was associated to MMPs, we used a broad-spectrum MMP inhibitor, GM6001. After GM6001 treatment (1 μM), the DQ-gelatin signal was significantly reduced (data not shown).

### *In situ* hybridization

Experimental and control chick embryos at stage HH11-15 were processed for *in situ* hybridization using *Timp2* probe. The *Timp2 in situ* hybridization probe was generated by PCR on cDNA from dissected region between the otic placode and the third somite at stage HH10-12. We used the primers 5’-ATGAGGCTTTCTGGGACGCG-3’ and 5’-TTTCCTACTGGCTACTGGAAT-3’ to PCR amplify the entire chicken *Timp2* gene (NM_204298). The amplified sequence was ligated into the PCRII vector using the TA cloning kit (Invitrogen, K2050-01). The plasmid was linearized with XhoI (Thermo Fisher Scientific ER0691), and the probe amplified with Sp6 polymerase (Promega), and digoxigenin (DIG)-labeled nucleotides (Roche). *In situ* hybridization was performed on wildtype and post-electroporation HH11-15 chicken embryos, as previously described (Acloque et al., 2008). After hybridization, the whole mount embryos were embedded in 20% sucrose (diluted in PBS). Cross sections with 12-18μm of optic region were collected by cryosectioning. We mounted the sections using FluoroShield Mounting (Abcam, United States). Images were acquired with Zeiss Axio Imager-D2 microscope coupled with an Axiocam 503 Color camera.

### Analysis of single-cell transcriptome data of mouse embryos at stage E9.5 and 10.5 from the Mouse Organogenesis Cell Atlas

We downloaded a single cell transcriptome dataset available at Mouse Organogenesis Cell Atlas (https://oncoscape.v3.sttrcancer.org/atlas.gs.washington.edu.mouse.rna/downloads) (Cao et al., 2019). These data were obtained from whole mouse embryos collected ranging from E9.5–13.5 (Cao et al., 2019). Cao et al profiled the RNA in nuclei using sci-RNA-seq3 and sequenced about 5000 raw reads per cell. For our analysis, we used the following available files: “gene_count_cleaned.RDS” (dgCMatrix with the count of 26183 genes and 1331984 cells from the 5 mouse embryos stages; cells were previously filtered and correspond to high-quality cells), ‘cell_annotate.csv’, ‘gene_annotate.csv’. To analyze the single-cell data we used RStudio and Seurat package (version 4.0.3) (Butler et al., 2018; Hao et al., 2021; Satija et al., 2015; Stuart et al., 2019).

## ACKNOWLEDGEMENTS

C.G.M. received CAPES and CNPq Graduate Fellowships. C.Y.I.Y. was funded by FAPESP grants 2017/07405-7, 2018/05958-1, 2020/07008-0. Additional funding includes NIH/NEI R01 EY012200 (A.C.). We would like to thank Marley J. dos Santos and Kveta Cveklova for superb technical and personal support. Prof. Estela Bevilacqua is acknowledged for providing the fluorescence microscope that was important to the progress of this work. The authors thank Michael Camerino, Jie Zhao and Dr. Danielle Rayêe for their help with mouse colonies breeding and maintenance. Dr. Danielle Rayêe and Dr. Mario Cruz are acknowledged for their crucial assistance in imaging mouse embryos in widefield and confocal fluorescence microscopy. We would like to thank SAG and JG and the rest of the Srinivas lab for the gift of some of the mouse tissue samples as well as constructive discussions.

