## Supplementary Figures for "Lens Placode Modulates Extracellular Matrix Formation During Early Eye Development"

**Supplemental Figure 1. Lamininα1 labelling becomes faulty in placodal region in mouse.** (A-D) Apical view from eye region of two mouse embryos in stage E9.0, phase 0 (A-B) and phase 1 (C-D) of lens placode development.. (A and C) Bottom optical slice (most basal optic section) of DAPI staining (grey), where we see the optic vesicle boundary that defines the optical region (dotted yellow line). (B and D) Z-projection of Lamininα1 (lama1, green) staining with lens pre-placode region (white arrowheads). (a-d) YZ orthogonal (optic slices) images. (a-b) The orthogonal view shows an intense Lamininα1 staining between the optic vesicle (OV) and pre-placodal ectoderm. DAPI staining (gray) shows the spherical nucleus inside the cuboidal pre-placodal cells. Lama1 (green) staining is intense in the entire optic slice, with no clear difference between the optic and non-optic domains. (c-d) The orthogonal view shows decreased Lama1 labelling between the optic vesicle and lens placode in phase 1, where the tissue is changing from cuboidal to columnar shape (white arrowhead). Between cuboidal placodal cells (phase 0) and the optic vesicle, Lama1 shows an intense labelling. Scale bar in A and C: 100 µm.

**Supplemental Figure 2. Lamininα1 labeling is more intense in the optic vesicle basal domain than lens placode during invagination in mouse.** (A-C) Confocal image of the lateral view from eye region of two mouse embryos in stage E11.0 (late phase 2b). Left column shows Lamininα1 staining, and the right column shows DAPI staining. (A) Z-projection of Lamininα1 (Lama1, gray) staining, shows a fibrillar staining pattern in non-placodal ectoderm (cyan arrowheads). While the lens placode invaginates the lens surrounding ectoderm approximates the invagination center (green arrowheads) to close the tissue and form the cornea. The ECM in this region has a fibrillar Lama1 pattern (cyan arrowheads). (B and C) Last optical slice (most basal optic section) of DAPI and Lama1 staining. The optic vesicle (OV) invaginates together with the lens placode invagination. (C) Higher magnification of image B shows the difference between Lama1 in the basal domain of the lens placode and the basal domain of the optic vesicle. Close to the optic vesicle, Lama1 staining is intense and fibrillar (magenta arrowhead), while close to the lens placode, it non-homogenous and less intense (white arrowhead).

**Supplemental Figure 3. Optic cells top differentially expressed genes.** (A) Expression levels of early optic vesicle, neuro retina, and retinal pigment epithelium markers in the UMAP graph. None of these optic vesicle markers are expressed in the lens placode cluster. (B) Table displaying the top 20 differentially expressed genes from each cluster. Avg_log2FC represents the average fold change values, and p_val_adj represents the adjusted p-value.

**Supplemental Figure 4. *In silico* analysis of mouse dissected eye scRNAseq data.** (A) Simplified workflow for scRNAseq data analysis. We used raw gene expression data from wildtype mouse dissected eyes at stage E9.5, from Yamada et al., 2021. The dataset is available at the NCBI Gene Expression Omnibus platform (accession number: GSE167126). The authors profiled the RNA in nuclei using Chromium Single Cell 3' Reagent Kits v3 (10x Genomics, Pleasanton, CA, USA) and sequenced with Illumina HiSeq 2500. For our analysis, we used only the data from wild-type animals from the files: 'GSM5097169_barcodes_WT26.tsv.gz', 'GSM5097169_genes_WT26.tsv.gz', GSM5097169_matrix_WT26.mtx.gz'. Similarly to the first scRNAseq analysis described here, we followed the workflow proposed by Satija Laboratory (available in https://satijalab.org/seurat/). From the total dataset of 1680 cells, we extracted the subset of only optic vesicle and ectodermal cells, resulting in 1025 cells. (B) UMAP graph displaying the clusters obtained from the analysis. (C) Expression levels of specific markers in different cell types: optic cells (*Pax6*, *Sox2*), non-neural ectoderm (*Cdh1*), lens placode (*Maf*, *Pitx3*), early optic vesicle (*Lhx2*, *Rax*), neuro retina (*Vsx2*), and retinal pigment epithelium (*Mitf*). (D) Dot plot graph showing the expression levels of ECM genes.

### SUPPLEMENTARY FIGURES

#### Supplemental Figure 1

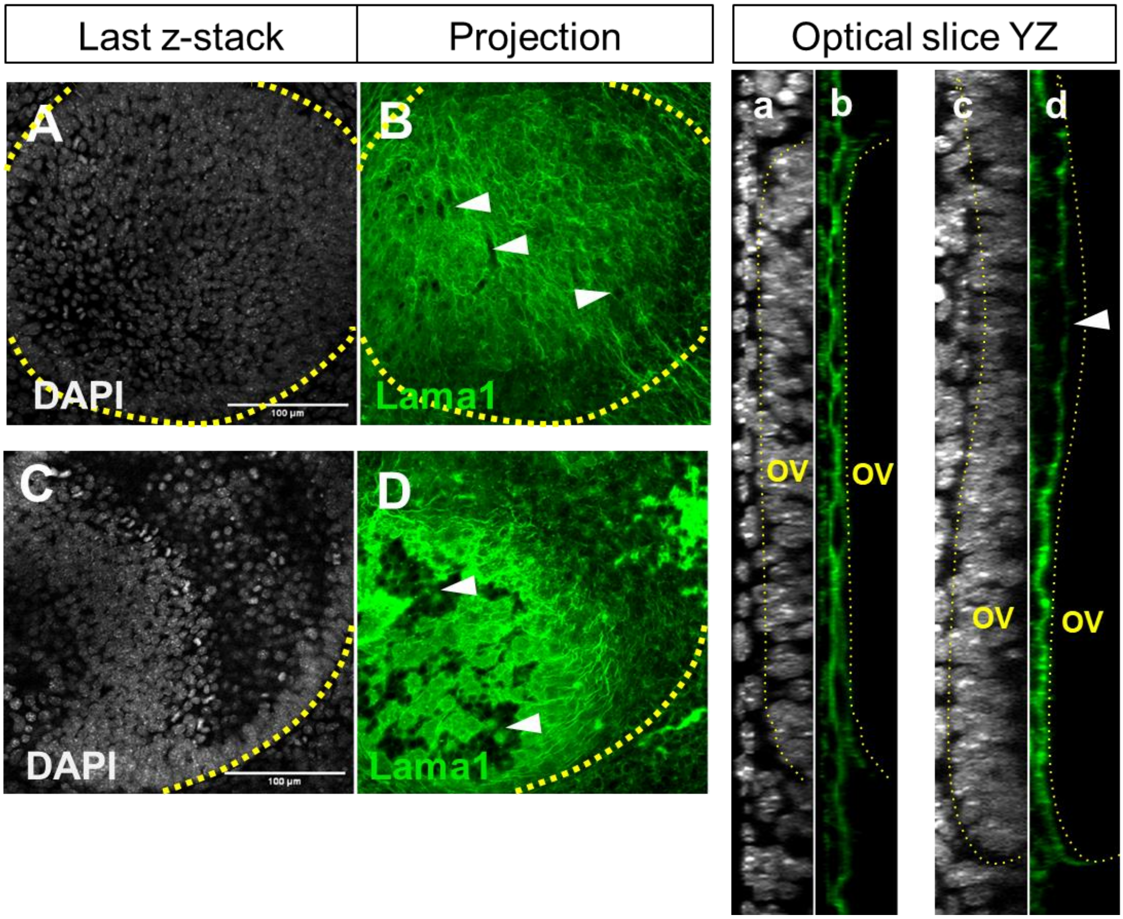

### Supplemental Figure 2

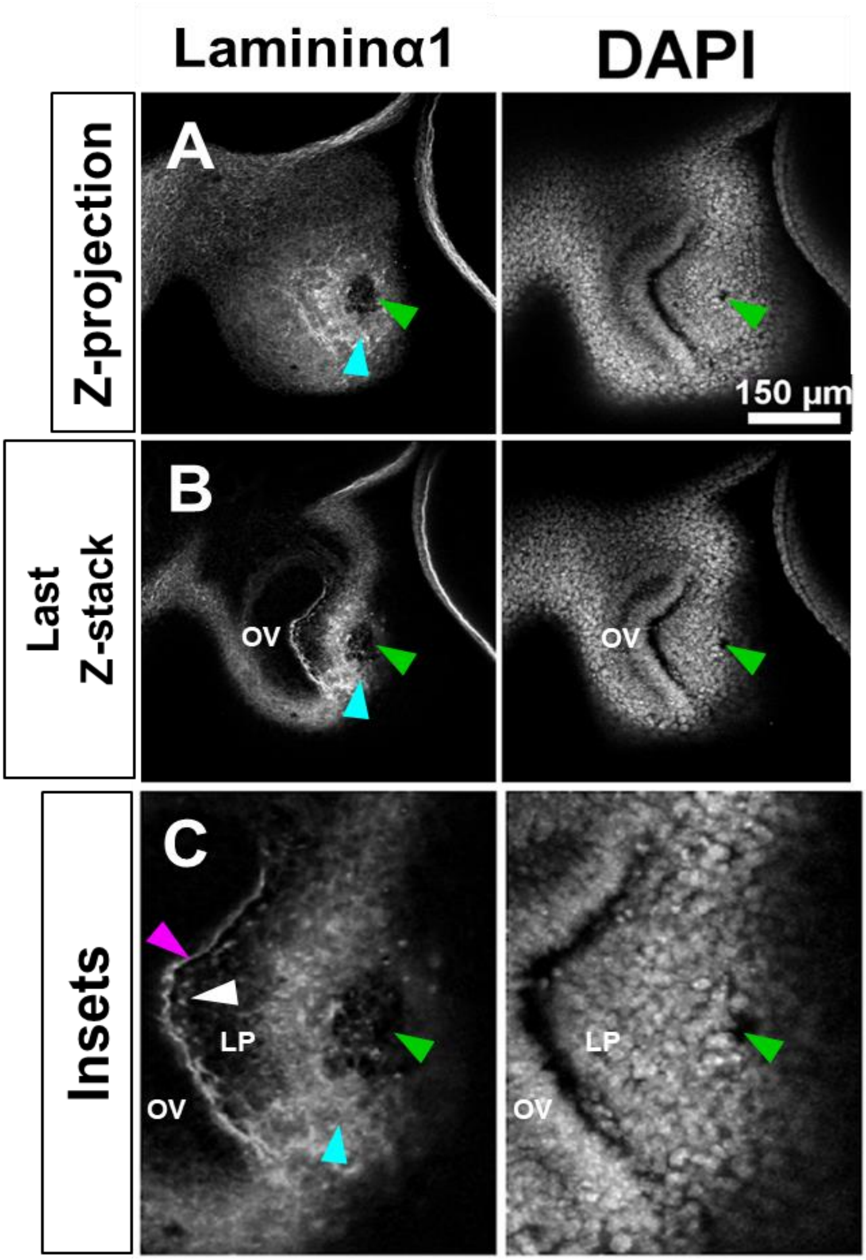

### Supplemental Figure 3

A

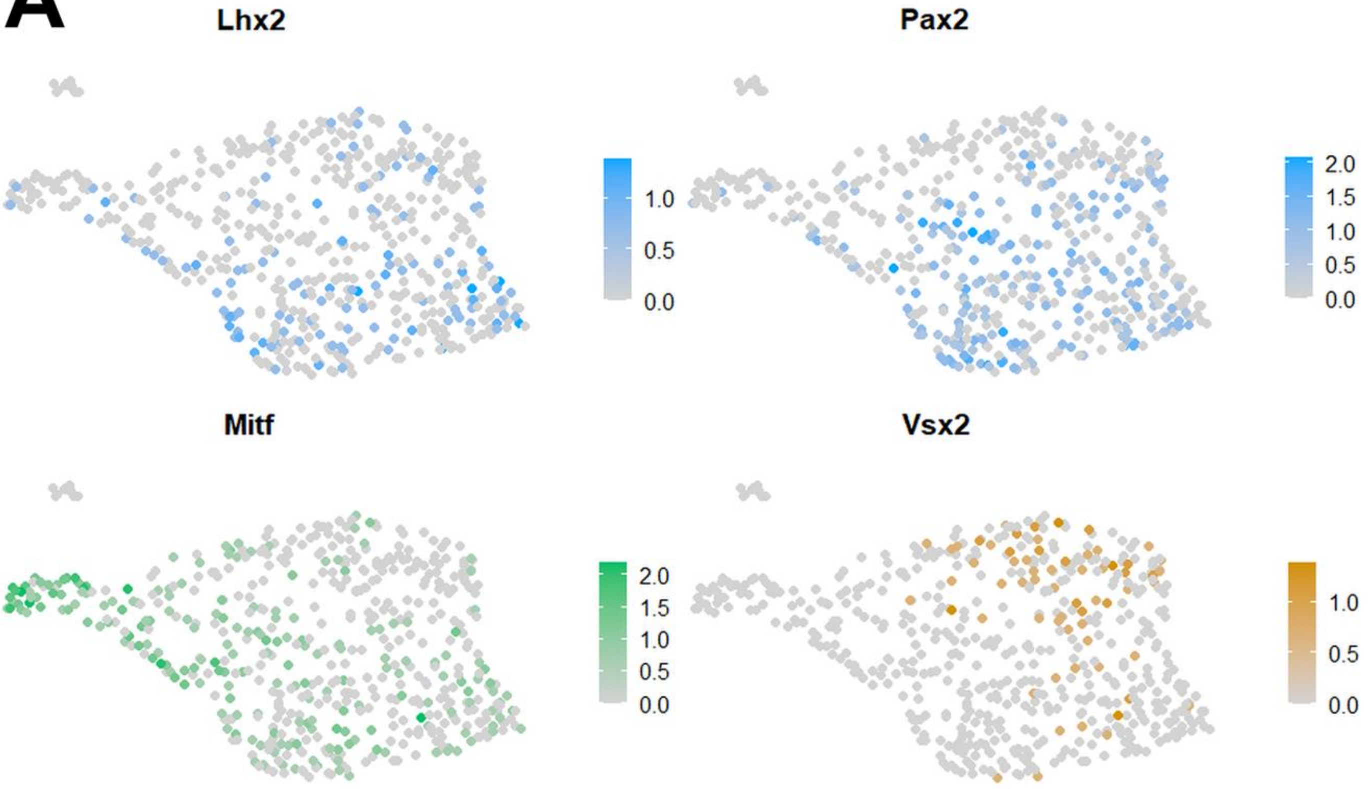


B

**LP**

| Gene | avg_log2FC | p_val_adj |
|---|---|---|
| Nid1 | 0,46 | 2,31E-08 |
| Tnc | 0,54 | 1,12E-07 |
| P3h2 | 1,77E+14 | 9,80E-04 |
| Cngb3 | 0,44 | 5,94E+01 |
| Ldlrad4 | 0,41 | 2,12E+02 |
| Ptprt | 0,86 | 6,20E+02 |
| Jazf1 | 0,68 | 7,07E+02 |
| Itga9 | 0,64 | 1,45E+04 |
| Igfbp7 | 0,72 | 3,66E+03 |
| Plxna4 | 0,72 | 2,38E+05 |
| Prox1 | 0,49 | 1,00E+06 |
| Airn | 1,48E+14 | 2,42E+07 |
| **Maf** | **0,64** | **2,50E+08** |
| Ccbe11 | 0,49 | 3,59E+08 |
| Col13a1 | 0,49 | 8,80E+07 |
| Pde4d | 1,90E+14 | 7,46E+09 |
| Etv1 | 0,41 | 1,53E-04 |
| Cadps2 | 0,48 | 4,94E-04 |
| **Sox2ot** | **0,84** | **8,39E-04** |
| Ell2 | 0,50 | 2,30E-03 |

**EarlyOV I**

| Gene | avg_log2FC | p_val_adj |
|---|---|---|
| Vwc2 | 0,45 | 6,56E+00 |
| Cdh20 | 0,61 | 1,92E+02 |
| Erbb4 | 0,74 | 9,44E+02 |
| Prtg | 0,61 | 1,27E+04 |
| Adgrl3 | 0,63 | 1,19E+05 |
| Efna5 | 0,48 | 1,95E+08 |
| Dach2 | 0,46 | 2,06E+08 |
| **Pax2** | **0,38** | **1,36E+09** |
| Grik2 | 0,40 | 4,46E+09 |
| Greb1l | 0,32 | 7,41E-04 |
| Cdh6 | 0,39 | 8,83E-04 |
| Nlgn1 | 0,40 | 3,79E-03 |
| Nrcam | 0,28 | 6,28E-03 |
| Pknox2 | 0,33 | 1,68E-02 |
| Plcxd3 | 0,32 | 1,99E-02 |
| Snd1 | 0,30 | 8,64E-02 |
| Rian | 0,32 | 1,25E-01 |
| P4hb | 0,26 | 1,87E-01 |
| Tenm4 | 0,36 | 4,97E-01 |
| Lsamp | 0,34 | 5,38E-01 |

**EarlyOV II**

| Gene | avg_log2FC | p_val_adj |
|---|---|---|
| Dtl | 0,61 | 4,88E+03 |
| Hells | 0,37 | 8,06E-03 |
| Hat1 | 0,32 | 9,66E-02 |
| Dip2b | 0,37 | 1,50E-01 |
| Neto2 | 0,34 | 4,71E-01 |
| Raly | 0,36 | 5,06E-01 |
| Lef1 | 0,31 | 5,38E-01 |
| E2f3 | 0,33 | 1,00E+00 |
| Scd2 | 0,43 | 1,00E+00 |
| Arih2 | 0,30 | 1,00E+00 |
| Zswim6 | 0,37 | 1,00E+00 |
| Usp6nl | 0,31 | 1,00E+00 |
| Eprs | 0,26 | 1,00E+00 |
| Atad5 | 0,30 | 1,00E+00 |
| Cdc73 | 0,26 | 1,00E+00 |
| Rab11fip3 | 0,26 | 1,00E+00 |
| Usp37 | 0,26 | 1,00E+00 |
| Rnf144a | 0,27 | 1,00E+00 |
| Cecr2 | 0,35 | 1,00E+00 |
| Zfp609 | 0,39 | 1,00E+00 |

**NR**

| Gene | avg_log2FC | p_val_adj |
|---|---|---|
| Cacna1d | 0,67 | 2,54E-10 |
| Epha6 | 0,93 | 1,38E-09 |
| Fbxl7 | 1,01E+14 | 3,57E-06 |
| Sfrp2 | 0,74 | 1,14E-03 |
| Cdh11 | 0,87 | 1,15E-03 |
| Sema3a | 0,74 | 1,23E-02 |
| Mecom | 0,75 | 7,17E-02 |
| Pcdh7 | 0,86 | 2,65E-01 |
| Ptprk | 0,90 | 8,67E-01 |
| Plekhg1 | 0,59 | 3,26E+00 |
| Ctnnd2 | 0,68 | 8,26E+00 |
| **Vsx2** | **0,40** | **2,43E+02** |
| Aldh1a1 | 0,60 | 2,63E+02 |
| Itgb8 | 0,46 | 2,33E+04 |
| Sema3e | 0,50 | 1,16E+05 |
| Celf2 | 0,57 | 8,08E+05 |
| Tox | 0,47 | 2,30E+06 |
| Sorcs1 | 0,71 | 2,21E+06 |
| Ltbp1 | 0,71 | 2,59E+07 |
| Cdon | 0,54 | 3,09E+07 |

**RPE I**

| Gene | avg_log2FC | p_val_adj |
|---|---|---|
| Robo2 | 1,16E+14 | 2,10E+04 |
| Tenm3 | 1,11E+14 | 9,91E+04 |
| Ralyl | 0,74 | 9,17E+07 |
| Aldh1a3 | 0,49 | 1,18E+08 |
| Hcn1 | 0,41 | 8,65E+08 |
| Pcdh9 | 0,77 | 5,00E+09 |
| Tenm2 | 0,57 | 2,39E-04 |
| Trpm31 | 0,57 | 3,15E-04 |
| 1700112E06Rik | 0,31 | 3,97E-04 |
| **Mitf** | **0,45** | **5,82E-04** |
| Pdzrn3 | 0,65 | 1,65E-03 |
| Rbms1 | 0,52 | 2,13E-03 |
| Gng12 | 0,27 | 9,01E-03 |
| Kcnj3 | 0,42 | 1,32E-02 |
| Mab21l2 | 0,25 | 3,52E-02 |
| Mir99ahg | 0,49 | 3,71E-02 |
| Tmem132c | 0,39 | 6,68E-02 |
| Pbx3 | 0,55 | 6,73E-02 |
| mt-Rnr21 | 0,56 | 7,81E-02 |
| Gpc61 | 0,36 | 1,65E-01 |

**RPE II**

| Gene | avg_log2FC | p_val_adj |
|---|---|---|
| Oca2 | 1,69E+14 | 4,16E-66 |
| Tyr | 2,04E+14 | 2,13E-63 |
| Gabrb3 | 2,71E+13 | 6,37E-46 |
| Il1rapl2 | 1,39E+14 | 6,39E-36 |
| Il1rapl1 | 2,13E+14 | 2,19E-34 |
| Slc4a5 | 0,97 | 2,02E-30 |
| Pmel | 0,88 | 2,91E-29 |
| Fstl4 | 1,61E+13 | 1,57E-27 |
| Gpnmb | 0,63 | 4,67E-22 |
| Dct | 1,04E+14 | 8,67E-21 |
| Mgll | 0,91 | 9,07E-20 |
| Col8a1 | 0,77 | 4,05E-18 |
| Bace2 | 0,48 | 4,62E-18 |
| Trpm1 | 1,41E+14 | 4,96E-17 |
| Col25a1 | 1,35E+14 | 4,82E-15 |
| Kcnip4 | 1,02E+14 | 2,42E-12 |
| Adcy8 | 0,78 | 1,26E-11 |
| 1700112E06Rik1 | 1,76E+14 | 1,92E-11 |
| Ankrd44 | 1,11E+14 | 7,62E-11 |
| Fam13a | 0,50 | 9,65E-10 |

### Supplemental Figure 4

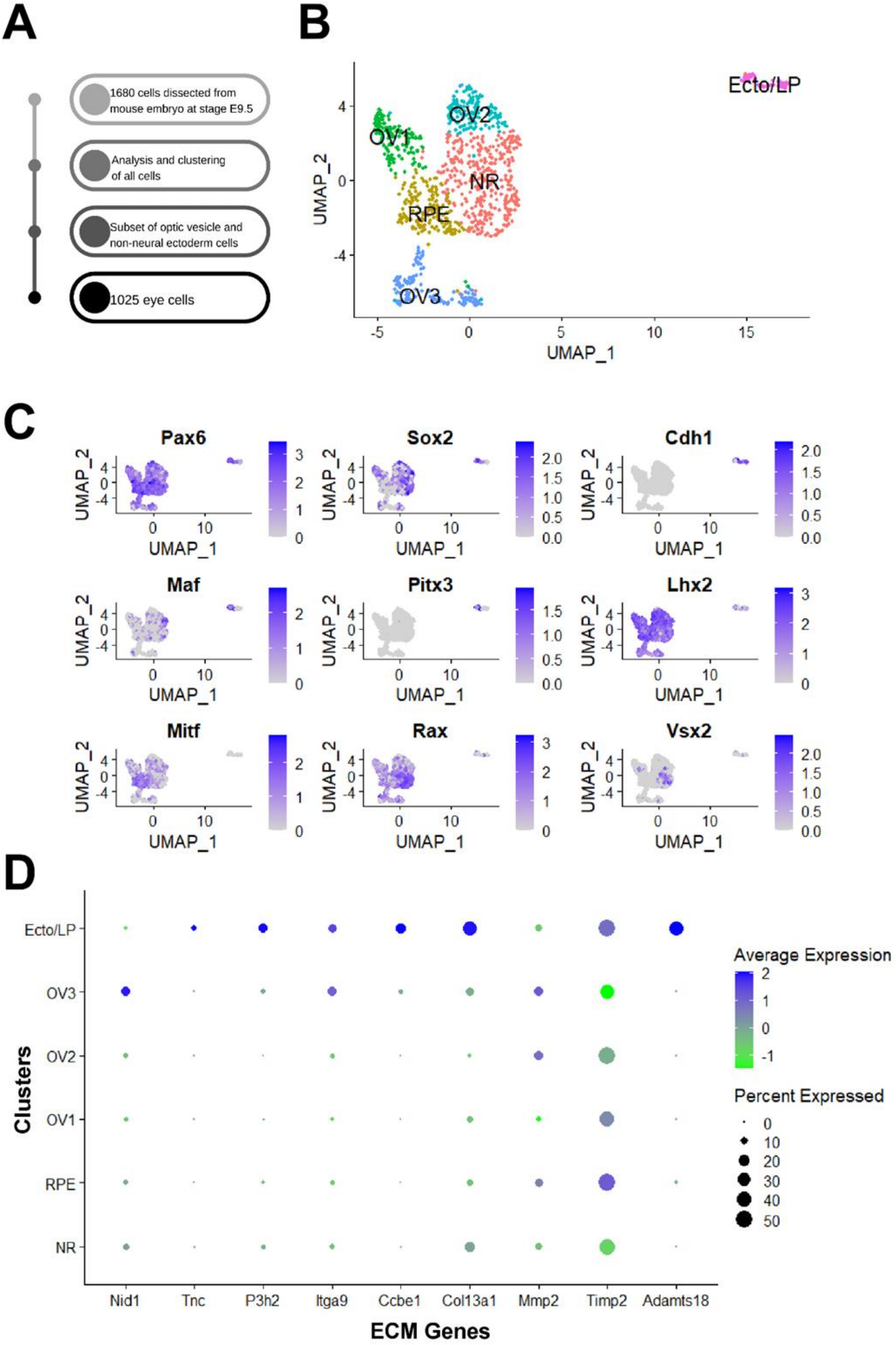
